# A fungal avirulence factor encoded in a highly plastic genomic region triggers partial resistance to septoria tritici blotch

**DOI:** 10.1101/264226

**Authors:** Lukas Meile, Daniel Croll, Patrick C. Brunner, Clémence Plissonneau, Fanny E. Hartmann, Bruce A. McDonald, Andrea Sánchez-Vallet

## Abstract

- Cultivar-strain specificity in the wheat-*Zymoseptoria tritici* pathosystem determines the infection outcome and is controlled by resistance genes on the host side, of which many have been identified. However, on the pathogen side, the molecular determinants of specificity are largely unknown.
- We used genetic mapping, targeted gene disruption and allele swapping to characterize the recognition of the new avirulence factor Avr3D1. We then combined population genetic and comparative genomic analyses to estimate the evolutionary trajectory of *Avr3D1*.
- Avr3D1 is specifically recognized by cultivars harboring the resistance gene Stb7 and triggers a strong defence response without preventing pathogen infection and reproduction. *Avr3D1* resides in a cluster of putative effector genes located in a region populated by independent transposable element insertions. The gene is present in all 132 investigated strains and is highly polymorphic, with a total of 30 different protein variants. We demonstrated that certain amino acid mutations in Avr3D1 led to evasion of recognition.
- These results demonstrate that quantitative resistance and gene-for-gene interactions are not mutually exclusive *per se*. Location of avirulence genes in highly plastic genomic regions likely facilitates accelerated evolution that enables escape from recognition by resistance proteins.

## Introduction

Whether mutualistic or parasitic, colonizing microbes evolve a high degree of specialization to recognize and infect their hosts and overcome host-inducible defences (van der Does & Rep, 2017). Host manipulation is frequently achieved by the secretion of effectors, which are often small secreted proteins (SSPs) that support growth and development of the microbe by conferring protection against host antimicrobial compounds or by altering host metabolism (Lo Presti *et al.,* 2015). Although effectors are beneficial for host colonization, some are specifically recognized by certain host genotypes, triggering an immune response (Jones & Dangl, 2006; Lo Presti *et al.,* 2015). This interaction typically follows the gene-for-gene model, in which a so-called resistance protein recognizes an effector, which is then called an avirulence factor (Flor, 1971; Jones & Dangl, 2006). A common assumption is that resistance/avirulence gene interactions confer complete resistance, whereas quantitative resistance, understood here as incomplete or partial resistance that allows some pathogen infection and reproduction, is based on different, race-nonspecific and therefore avirulence-independent mechanisms. This paradigm originated from work on biotrophic pathogens, where avirulence recognition often leads to complete immunity via induction of a hypersensitive response (Cook *et al.,* 2015; Niks *et al.,* 2015). But it is often overlooked that gene-for-gene interactions could also lead to quantitative resistance, as suggested by several studies (Antonovics *et al.,* 2011; Rietman *et al.,* 2012; Chen *et al.,* 2013). Recently, more refined concepts such as the “invasion model” (Cook *et al.,* 2015) or "effector-triggered defence” (Stotz *et al.,* 2014) emphasized a broader perspective for the gene-for-gene model, in which resistance gene-based effector recognition and quantitative resistance are not mutually exclusive (Niks *et al.,* 2015). However, avirulence factors leading to quantitative resistance have only rarely been described (Schirawski *et al.,* 2010; Rietman *et al.,* 2012).

Host recognition of effectors exerts an evolutionary pressure that favors sequence modification, deletion or acquisition of new effectors to overcome the immune response. Thus, genes encoding effectors are among the most polymorphic found in pathogen genomes (Win *et al.,* 2012). The mechanisms underlying effector diversification remain largely unexplored. Many pathogen genomes are compartmentalized into highly conserved or rapidly evolving regions, often described as the “two-speed genome” (Raffaele & Kamoun, 2012). Effector genes are frequently localized in the highly variable compartments, which are often rich in transposable elements (Ma *et al.,* 2010; Soyer *et al.,* 2014; Plissonneau *et al.,* 2018). Transposable elements are thought to contribute to genome evolution and the diversification of effector genes (Raffaele & Kamoun, 2012). They translocate within a genome causing gene disruption, duplication or deletion of genomic sequences. In addition, transposable elements contribute to variability by favoring non-homologous recombination or through repeat-induced point mutations (RIP) (Möller & Stukenbrock, 2017; Seidl & Thomma, 2017). Pathogens carrying these highly plastic genome regions are thought to benefit from an increased versatility to adapt to different conditions or to an evolving host (Dong *et al.,* 2015; Faino *et al.,* 2016).

The most damaging pathogen of wheat in Europe is *Zymoseptoria tritici,* an ascomycete fungus that causes septoria tritici blotch (Fones & Gurr, 2015). Fungal hyphae penetrate the stomata and colonize the apoplast in a long asymptomatic phase that lasts between 7 and 14 days, depending on the weather conditions, the host cultivar and the pathogen strain. This period is followed by a rapid induction of necrosis that is accompanied by the development of asexual reproductive structures called pycnidia, which contain asexual spores that spread the disease during a growing season (Kema *et al.,* 1996; Duncan & Howard, 2000). The genetic basis of *Z. tritici* virulence is poorly understood as a result of its largely quantitative nature (Hartmann *et al.,* 2017; Stewart *et al.,* 2018). Two highly conserved LysM effectors, Mg1LysM and Mg3LysM, prevent fungal recognition and shield the fungal cell wall from degradation by host hydrolytic enzymes (Marshall *et al.,* 2011). The other known effectors of *Z. tritici,* Zt80707, AvrStb6 and Zt_8_609, are rapidly evolving small secreted proteins (Poppe *et al.,* 2015; Hartmann *et al.,* 2017; Zhong *et al.,* 2017). The latter two were identified because they are specifically recognized by certain wheat cultivars, and they were found to be located in transposable element-rich genomic regions (Brading *et al.,* 2002; Hartmann *et al.,* 2017; Zhong *et al.,* 2017). AvrStb6 is recognized by the resistance protein Stb6 in a gene-for-gene interaction that leads to a strong resistance response, completely blocking the progression of the infection (Kema, GH *et al.,* 2000; Brading *et al.,* 2002). In addition to *Stb6,* 19 other race-specific *Stb* resistance genes with large effects have been mapped, but their corresponding avirulence factors remain unknown (Brown *et al.,* 2015). We hypothesized that one of these *Stb* genes might be responsible for the differences in resistance of cultivar Runal to two Swiss strains (3D1 and 3D7) (Stewart *et al.,* 2018). The more virulent strain 3D7 produced necrotic lesions faster than 3D1. The less virulent 3D1 strain was successful in producing pycnidia, but at a lower density and with a less uniform distribution across the leaf surface than 3D7 (Fig. S1). A single, large-effect quantitative trait locus (QTL) encoding differences in lesion size and pycnidia density between 3D1 and 3D7 was mapped to a region on chromosome 7 (Stewart *et al.,* 2018). However, the genes responsible for the differences in virulence were not identified.

Here we aimed to broaden our knowledge of the genetic basis of host-race specificity in *Z. tritici.* First, we showed that *Avr3D1* is the gene responsible for the differences in quantitative virulence between 3D1 and 3D7. We then demonstrated that Avr3D1 is an avirulence factor whose recognition is host-specific, but triggers an incomplete, quantitative resistance. We next studied the evolutionary trajectory of *Avr3D1* by combining population genetic and comparative genomic analyses involving 132 *Z. tritici* strains originating from four field populations on three continents as well as 11 strains of the closest known relatives of *Z. tritici.* We found that *Avr3D1* is a member of an effector gene cluster that is located in a highly dynamic genomic region containing many independent insertions involving different families of transposable elements. Because an intact and presumably functional version of *Avr3D1* was found in all strains of *Z. tritici* and in its closest relatives, we conclude that *Avr3D1* plays an important role in the life history of *Z. tritici.* Maintaining *Avr3D1* in a highly plastic genomic region likely provides an advantage by accelerating evolution that enables an escape from recognition in wheat populations carrying the corresponding resistance gene.

## Materials and methods

### QTL mapping

To generate a genetic map, we used the previously generated RADseq data from the progeny of the cross between 3D7 and 3D1 (Lendenmann *et al.,* 2014). Quality trimmed reads were aligned to the genome of 3D7 (Plissonneau *et al.,* 2016) using bowtie2 with default parameters (Langmead & Salzberg, 2012). SNPs were called in each progeny with the HaplotypeCaller tool from GATK v3.3 (McKenna *et al.,* 2010) and further filtered for their quality using the following parameters: > QUAL 5000, QD > 5, MQ > 20, and ReadPosRankSum, MQRankSum, and BaseQRankSum between ×2 and 2. We constructed the linkage map using R/qtl v1.40-8 (Arends *et al.,* 2010). We retained only progenies for which 45% of all SNPs were genotyped, then we removed SNPs genotyped in less than 70% of the progenies. Potential clones (i.e. progenies with more than 90% shared SNPs) were excluded. We removed adjacent nonrecombining markers. QTL mapping was performed with the QTL package in R (R-Core-Team, 2013), similar to the procedure described by (Lendenmann *et al.,* 2014) using the pycnidia density dataset (Stewart *et al.,* 2018).

### *Z. tritici* and bacterial strains

The Swiss strains ST99CH_3D1 (3D1) and ST99CH_3D7 (3D7) (described in (Linde *et al.,* 2002)) or mutant lines derived from them were used in this study. Standard conditions for *Z. tritici* cultivation consisted of yeast-sucrose broth (YSB) medium (10 g/L yeast extract, 10 g/L sucrose, 50 μg/ml kanamycin sulfate) at 18°C or yeast-malt-sucrose (YMS) medium (4 g/L yeast extract, 4 g/L malt extract, 4g/L sucrose, 12 g/L agar) at 18°C. For molecular cloning and plasmid propagation, *E. coli* strains HST08 (Takara Bio, USA) or NEB^®^ 5-alpha (New England Biolabs) were used. *Agrobacterium tumefaciens-mediated* transformation was performed with *A. tumefaciens* strain AGL1. If not stated otherwise, *E. coli* and *Agrobacterium* lines were grown in Luria Bertani (LB) medium containing kanamycin sulfate (50 μg/mL) at 37°C or in LB medium containing kanamycin sulfate (50 μg/mL), carbenicillin (100 μg/mL) and rifampicin at 28°C (50 μg/mL), respectively.

### Generation of plasmid constructs for targeted gene disruption and ectopic gene integration

All PCR reactions for cloning procedures were performed using NEB^®^ Phusion polymerase (New England Biolabs) with primers listed in Table S1. All DNA assembly steps were conducted with the In-Fusion^®^ HD Cloning Kit (Takara Bio,USA) following the manufacturer’s instructions. To create constructs for targeted gene disruption, two flanking regions of at least 1 kb in size for homologous recombination were amplified from *Z. tritici* genomic DNA. The hygromycin resistance gene cassette, used as a selectable marker, was amplified from pES6 (Eva H. Stukenbrock, unpublished). The three fragments were assembled into pES1 (Eva H. Stukenbrock, unpublished) and linearized with KpnI and PstI (New England Biolabs), resulting in pES1Δ581_3D1_ and pES1Δ581_3D7_. To create the construct for ectopic integration of *Avr3D1_3D1_,* a fragment containing *Avr3D1_3D1_* including the 1.3kb sequence upstream of the start codon and the 1-kb sequence downstream of the stop codon was amplified and cloned into pCGEN (Motteram *et al.,* 2011) that had been linearized with KpnI, resulting in pCGEN-581_3D1_ect. To exchange the CDS in pCGEN-581_3D1_ect, we first digested it with XhoI (New England Biolabs) to linearize it and remove the *Avr3D13D1* CDS. In this digestion, the promoter was partially removed from the vector. In a second step a fragment containing the CDS and intron 1 of *Avr3D1_3D7_* (amplified from 3D7 genomic DNA) and a fragment to reconstitute the promoter sequence of *Avr3D13D1* (amplified from pCGEN-581_3D1_ect) were assembled into the linearized pCGEN-5813D1ect, resulting in pCGEN-5813D7ect. Constructs were transformed into *E. coli* by heat shock transformation, mini-prepped and verified by diagnostic digests and Sanger sequencing (MicroSynth, Switzerland). Confirmed plasmids were transformed to *A. tumefaciens* cells by electroporation.

### Agrobacterium tumefaciens-mediated transformation (ATMT) of Zymoseptoria tritici cells

ATMT of *Z. tritici* was performed according to Zwiers & De Waard, 2001 with the following modifications: *A. tumefaciens* lines were grown as liquid cultures for approx. 24 hrs. Cell concentrations were estimated by measuring the optical density (OD_600_) and the cultures were diluted to an OD_600_ of 0.15 in induction medium (pH 5.7, 50 μg/mL kanamycin sulfate, 100 μg/mL carbenicillin, 50 μg/mL rifampicin, 10 mM glucose, 200 μ M acetosyringone). These cultures were incubated at 28°C until they reached an OD_600_ of 0.25-0.35 and 100 μL were mixed with 100 μL of *Z. tritici* cell suspensions (cells grown on YMS for 4-6 days and washed off with water) and plated on induction medium covered with nitrocellulose membranes. After 3 days of incubation at 18°C, the nitrocellulose membranes were placed on YMS medium containing cefotaxime (200 μg/mL) and either hygromycin B (100 μg/mL) or geneticin (150 μg/mL), depending on the resistance cassette of the construct, and incubated at 18°C until colonies appeared. Colonies were streak-plated on the same selective medium to isolate single colonies before the mutant lines were grown on YMS without selection. For knockout lines, disruption of the target genes was verified using a PCR-based approach. We determined the copy number of the transgene by qPCR on genomic DNA extracted with DNeasy Plant Mini Kit (Qiagen). The target used was the selection marker and the reference gene was *TFC1* (Table S1). Only single insertion lines were selected for further experiments.

### Infection assays

Seeds from cultivars “Runal”, “Titlis”, “Drifter”, “Chinese Spring” and “Arina” were purchased from DSP Ltd. (Delley, Switzerland). Seeds were sown in peat substrate Jiffy^®^ GO PP7 (Jiffy Products International) and grown for 17 days in a greenhouse at 18°C (day) and 15°C (night) with a 16-hrs photoperiod and 70% humidity. For all infection experiments, square pots (11×11×12 ME, Lamprecht-Verpackungen GmbH, Germany) containing 16-18 seedlings or 2×3 pot arrays (7×7 cm and 200 mL each, Bachmann Plantec AG, Switzerland) containing two seedlings per unit were used. The infection procedure for the two pot types was identical. *Z. tritici* inoculum was prepared as follows: 50 mL of YSB medium were inoculated in 100-mL Erlenmeyer flasks from *Z. tritici* glycerol stocks stored at ×80°C. After 4-6 days of incubation (18°C, shaking at 120 rpm), liquid cultures were filtered through sterile cheesecloth and pelleted (3273 g, 15 min, 4°C). The supernatant was discarded and the cells were resuspended in sterile deionized water and stored on ice until infection (0-2 days). The concentrations of the spore suspensions were determined using KOVA^®^ Glasstic^®^ counting chambers (Hycor Biomedical, Inc., USA) and adjusted to 10^6^ spores/mL in 0.1% (v/v) Tween^®^ 20. Spore viability and concentration was analyzed by performing a developmental assay on YMS medium as described below. Plants were sprayed until run-off with 15 mL spore suspension per pot/array. Square pots were placed in plastic bags (PE-LD, 380×240 mm) to support the leaves and stems. Subsequently, they were placed in a second plastic bag (PE-LD 650×400 mm, two pots each), which was sealed to keep humidity at 100%. Pot arrays were placed directly into the sealing bags. After three days, the sealing bags were trimmed to a height of around 27 cm and then opened, in the case of the 2×3 pot arrays, or completely removed in the case of the square pots, leaving the supporting bags intact in the latter case. For symptom quantification, second or third leaves were mounted on paper sheets, scanned with a flatbed scanner (CanoScan LiDE 220) and analyzed using automated image analysis (Stewart *et al.,* 2016). Data analysis and plotting was performed using RStudio Version 1.0.143. Confidence intervals of the medians were determined using the “boot” package and Kolmogorov-Smirnov (KS) tests for statistical significance with the “Matching” package.

### RNA isolation and quantitative RT-PCR

Second leaves from cv. Runal were infected with 3D1 or 3D7, harvested and scanned as described. Immediately after scanning, the tip 2 cm of the leaves were excised and discarded and the adjacent 8.5 cm sections were frozen in N_2_. Three biological replicates were harvested. Leaf tissue was homogenized using a Bead Ruptor with a cooling unit (Omni International) and zirconium oxide beads (1.4 mm). RNA was isolated using the GENEzol reagent (Geneaid Biotech) and purified with the RNeasy Mini kit (Quiagen) including an on-column DNase treatment with the RNase-Free DNase Set (Qiagen) according to the manufacturer’s instructions. cDNA was produced with the RevertAid First Strand cDNA Synthesis Kit (Invitrogen), using up to 900 ng RNA (estimated with NanoDrop) per reaction. To determine expression of *Avr3D1* relative to the *18S* reference gene, qRT-PCR was performed with a LightCycler^®^ 480 (Roche) using white 384-well plates. Each reaction consisted of 250 nM of each primer, template cDNA generated from 11-30 ng of RNA and 1x HOT FIRE Pol^®^ EvaGreen^®^ qPCR Mix Plus mastermix (Solis BioDyne) in a total volume of 10 μL. Amplification was performed with a 10-min step of initial denaturation and enzyme activation and 40 cycles of 95°C (15 s) and 60°C (60 s). Each sample was run in technical triplicates. Relative expression was calculated with LightCycler^®^ 480 software using the “advanced relative quantification” tool. The mean and confidence interval of the mean was calculated with RStudio Version 1.0.143.

### Stress and development assay

The obtained *Z. tritici* mutant lines were tested for an altered, plant-unrelated phenotype under various conditions including stress by growing them on PDA, YMS and YMS supplemented with H2O2 (2 mM for 3D1 lines and 1 mM for 3D7 lines) or 1 M NaCl at 18°C. All media contained kanamycin sulfate (50 μg/mL). An additional stress condition consisted of growth at 28°C on YMS. Inoculum preparation and quantification were the same as for the infection assays. 2.5-μL drops of spore suspensions of 10^7^, 10^6^, 10^5^ and 10^4^ spores/mL were plated on the media described above. Plates were assessed after 6 days of upside-down incubation. Mutant lines exhibiting abnormal development or growth deficiencies were excluded from further experiments.

### Manual annotation of three small secreted proteins in the QTL for virulence

We used RNAseq raw data of IPO323 infecting wheat seedlings (Rudd *et al.,* 2015) to manually annotate the gene Zt09_7_00581. To annotate the other genes in the cluster, we used RNAseq raw data of 3D7 from two different experiments and at 6 different time points (Palma-Guerrero *et al.,* 2016). The data was previously deposited in NCBI with the experiment numbers SRP061444 and ERP009837. RNAseq reads were analysed as described in Hartmann & Croll 2017). Possible reading frames were manually examined using Integrative Genomics Viewer (IGV, Broad Institute, Robinson *et al.,* 2011). Signal peptides were predicted using Signal P 4.1 (CBS, Petersen *et al.,* 2011).

### *Zymoseptoria tritici* strain collections

We used 132 strains collected in four different countries (Switzerland, Israel, US and Australia; (Zhan *et al.,* 2005)). Whole-genome Illumina sequencing data of the 132 strains was previously deposited on the NCBI Short Read Archive under the BioProject ID numbers PRJNA178194 and PRJNA327615 (Torriani *et al.,* 2011; Croll *et al.,* 2013; Hartmann & Croll, 2017; Hartmann *et al.,* 2017). We used complete genome assemblies of IPO323, ST99CH_3D1 (3D1), ST99CH_3D7 (3D7), ST99CH_1E4 (1E4) and ST99CH_1A5 (1A5) previously described by Goodwin *et al.,* (2011) and Plissonneau *et al.,* (2016; 2018). BLAST searches were performed using the blastn command of the ncbi-blast-2.2.30+ software (Camacho *et al.,* 2009). Synteny of the QTL between IPO323, 3D1, 3D7, 1E4 and 1A5 was analyzed using blastn and visualized using the R package genoPlotR v. 0.8.4 (Guy *et al.,* 2010). Homologs of *Avr3D1* were identified by Blastn using CLC Genomic Workbench 9 (Qiagen) in the strains 3D1,3D7, 1E4 and 1A5.

We searched for orthologs of *Avr3D1* using the blast algorithm implemented in CLC Genomics Workbench 9 (Qiagen) in one strain of *Zymoseptoria passerinii* [NCBI genome accession no. AFIY01 (fungal strain SP63)], four strains of *Z. ardabiliae* [AFIU01 (STIR04 1.1.1), AFIV01 (STIR04 1.1.2), AFIW01 (STIR04 3.13.1), AFIX01 (STIR04 3.3.2)], one strain of *Z. brevis* Zb18110 (LAFY01), and five strains of the sister species Z. *pseudotritici* [AFIQ01 (STIR04 2.2.1), AFIO01 (STIR04 3.11.1), AFIR01 (STIR04 4.3.1), AFIS01 (STIR04 5.3), AFIT01 (STIR04 5.9.1)]. The genomes were downloaded from NCBI under the accession numbers PRJNA63035, PRJNA277173, PRJNA63037, PRJNA63039, PRJNA343335, PRJNA343334, PRJNA343333, PRJNA343332, PRJNA63049, PRJNA273516 and PRJNA46489.

### Presence/absence polymorphism of TEs and annotation

Repetitive DNA was identified for the 132 strains. For 3D1, 3D7, 1E4 and 1A5 full genome annotations were already available (Plissonneau *et al.,* 2016; Plissonneau *et al.,* 2018). We annotated and masked repetitive elements in the genomes of the remaining 128 strains using RepeatModeler version 1.0.8, as described before (Plissonneau *et al.,* 2016) and we masked the genomes using RepeatMasker version 4.0.5 with the library previously obtained for *Z. tritici* strain IPO323 (Grandaubert *et al.,* 2015) according to TE nomenclature defined by Wicker *et al.,* (2007).

### DNA and protein alignments and phylogenetic tree

DNA and protein sequence alignments of Avr3D1 and the other SSPs from different strains were obtained using CLC Genomics Workbench 9 (Qiagen). For the phylogenetic analysis amino acid sequence of Avr3D1 were aligned using Muscle. The Maximum Likelihood Phylogeny Reconstruction was performed applying WAG model, with the software Mega6 (Tamura *et al.,* 2013).

### Population genetic analysis

DnaSP v5 (Librado & Rozas, 2009) was used to calculate summary statistics of population genetic parameters associated with *Avr3D1.* Sliding window analyses of π were conducted using DnaSP with a window length set to 20 bp and a step size of 5 bp. The haplotype alignment of the coding region was used to generate a parsimony haplotype network using the TCS method (Clement *et al.,* 2000) as implemented in the PopART package v. 1.7 (Leigh & Bryant, 2015). TCS utilizes statistical parsimony methods to infer unrooted cladograms based on Templeton’s 95% parsimony connection limit. Mutational steps resulting in nonsynonymous changes were identified with DnaSP.

The degree of selection was estimated by comparing dN (the number of nonsynonymous changes per nonsynonymous site) with dS (the number of synonymous changes per synonymous site) for all pairwise sequence comparisons using DnaSP. A dN/dS ratio of 1 (ω = 1) indicates neutrality, while ω < 1 suggests purifying, and ω > 1 suggests diversifying selection. Since diversifying selection is unlikely to affect all nucleotides in a gene, ω averaged over all sites is rarely > 1. We focused on detecting positive selection that affects only specific codons in *Avr3D1* by applying the maximum-likelihood method CodeML implemented in the PAML software (Phylogenetic Analysis by Maximum Likelihood; (Yang, 1997; Yang, 2007)).

## Results

### Differences in virulence map to an effector gene cluster on chromosome 7

To identify the gene(s) responsible for the differences in virulence between 3D1 and 3D7, we generated a new linkage map based on the completely assembled genome of the parental strain 3D7 (Plissonneau *et al.,* 2016). Mapping onto the new genome sequence provided twice as many SNP markers and enabled the identification of additional crossovers that allowed us to reduce the number of candidate genes in the previously identified virulence QTL on chromosome 7 (Stewart *et al.,* 2018). The new map yielded a narrower QTL interval (LOD=41.5, p<10^−15^) located within the original QTL interval. The 95% confidence interval for the new QTL in 3D7 spanned 75 kb and contained only 4 of the 35 genes identified in the original QTL, including *Mycgr3T105313, Zt09_7_00581*, *Mycgr3T94659 (Zt09_7_00582)* and the predicted SSP-encoding gene *QTL7_5.* A manual RNAseq-supported reannotation in 3D7 of the confidence interval revealed two additional genes predicted to encode SSPs, which were named *SSP_3* and *SSP_4* (Fig. S2, Table S2). *Zt09_7_00581* was reannotated as also encoding a predicted SSP after identifying an upstream start codon (Fig. S2, Table S2). The four genes predicted to encode SSPs in 3D7 formed a cluster of putative effectors.

### Avr3D1 recognition contributes to quantitative resistance

In contrast to *SSP_3* and *SSP_4,* the genes *Zt09_7_00581* and *QTL7_5* are highly expressed during infection (Stewart *et al.,* 2018, Fig. S2b). Therefore, we considered them as the best candidate genes to explain the virulence QTL and they were selected for functional validation. Knockout mutants in both parental strains were generated by targeted gene disruption and used for virulence assessments in cv. Runal. Mutants in *QTL7_5* in the 3D7 and 3D1 backgrounds (3D7Δqtl7_5 and 3D1Δqtl7_5) did not show an altered phenotype when they were scored for host damage (Fig. S3), suggesting that *QTL7_5* is not involved in virulence on cv. Runal. Similarly, the virulence phenotype of the *Zt09_7_00581* mutant in the 3D7 background (3D7Δavr3D1) was unaltered compared to the wild type (Fig. 1A, S4), but disrupting *Zt09_7_00581* in 3D1 (3D1Δavr3D1) led to faster development of necrotic lesions and to the production of more pycnidia compared to the wild type 3D1 (Fig. 1A, S4, S5). Phenotypic alterations of the knockout lines in 3D1 were specific to *in planta* conditions, as no developmental alterations were observed when the mutants were grown on solid media used for stress assays (Fig. S6). The facts that *Zt09_7_00581* negatively affects virulence in 3D1 but not in 3D7 and that *in vitro* growth is unaffected by gene deletion suggests that this gene encodes an avirulence factor, so we renamed this gene *Avr3D1.* Even though Avr3D1 hinders the progression of the infection by 3D1, the avirulent strain is able to infect and produce pycnidia. Thus, Avr3D1 triggers a quantitative resistance response.

**Fig. 1.**
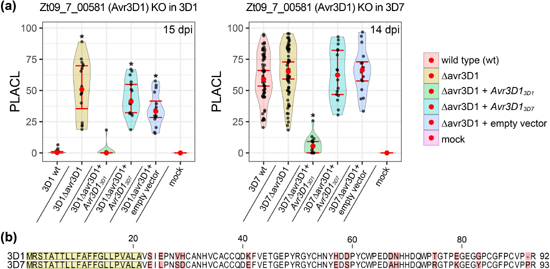
Z09_7_00581 encodes the avirulence factor Avr3D1. (a) Percentage of leaf area covered by lesions (PLACL) produced by the wildtype, the Avr3D1 knockout (KO, Δavr3D1) and the ectopic mutants expressing the Avr3D1 allele of either 3D1 (*Avr3D1*_3D1_) or 3D7 (*Δvr3D1*_3D7_) in the knockout background. Left panel: Mutants in the 3D1 background at 15 dpi. Right panel: Mutants in the 3D7 background at 14 dpi. Red dots represent the median of at least 15 leaves (except for the mock treatment, for which at least eight leaves were used), error bars represent 95% confidence intervals of the medians and black dots represent individual data points. Asterisks indicate statistical differences between wild type and knockout (p-value < 0.01, KS-test). (b) Amino acid sequence alignment of Avr3D1 variants of 3D1 and 3D7. The signal peptide sequence is highlighted in yellow and sequence polymorphisms between both alleles are shown in red.

To find out if 3D7 modulates the expression of *Avr3D1* to escape recognition, we quantified expression levels during infection for both strains. The expression pattern of *Avr3D1* in the virulent 3D7 strain was similar to 3D1, demonstrating that 3D7 is able to infect despite highly expressing *Avr3D1. Avr3D1* expression was high during the entire asymptomatic phase, peaking before the switch to the necrotrophic phase but dropping rapidly after the first symptoms appeared (Fig. S7), indicating a role for this SSP in host colonization, possibly during the asymptomatic phase, the switch to necrotrophy, or both.

### Avr3D1 is recognized by different wheat cultivars harboring *Stb7*

To determine if recognition of Avr3D1_3D1_ is mediated by a specific resistance protein, a set of 16 additional wheat cultivars was assessed for resistance against 3D1 and 3D1Aavr3D1. Three (Estanzuela Federal, Kavkaz-K4500 L.6.A.4 and TE-9111) out of 16 cultivars exhibited a significantly lower level of resistance against 3D1Δavr3D1 compared to 3D1 (Fig. 2, S8), suggesting the presence of a host-specific factor contributing to resistance against 3D1, possibly a resistance protein. In none of these three cultivars did the presence of Avr3D1 completely abolish lesion development and pycnidia production, demonstrating that the quantitative nature of Avr3D1_3D1_-induced resistance is a general phenomenon and not restricted to cv. Runal. All three cultivars that exhibited Avr3D1_3D1_-induced resistance were reported to carry the resistance gene *Stb7* (Brown *et al.,* 2015) and are also likely to carry the linked resistance gene *Stb12* (Chartrain *et al.,* 2005), leading us to propose Stb7 and Stb12 as candidate resistance proteins recognizing Avr3D13D1.

**Fig. 2.**
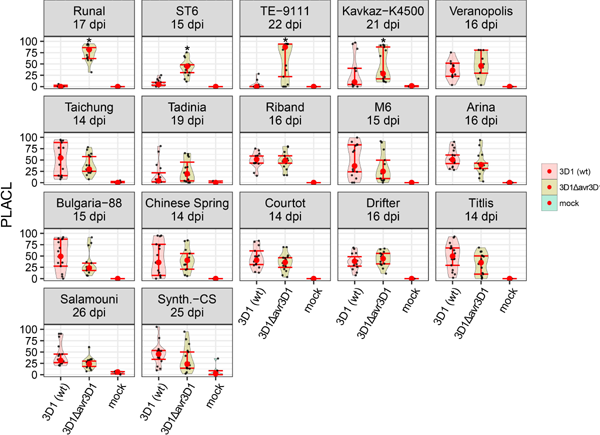
Avr3D1 is specifically recognized by four wheat varieties. Violin plots showing the percentage of leaf area covered by lesions (PLACL) produced by the wildtype 3D1, the Avr3D1 knockout (3D1Aavr3D1) and the mock control in seventeen wheat varieties. Harvesting time points varied because of cultivar-specific infection dynamics. Red dots represent the median of at least 10 leaves (except for the mock treatments, for which at least 4 leaves were used), error bars represent 95% confidence intervals of the medians and black dots represent individual data points. Asterisks indicate statistical differences between wild type and knockout (p-value < 0.01, KS-test). Synth. CS = Synthetic Chinese Spring; ST6 = Estanzuela Federal; M6 = M6 synthetic (W-7984), Kavkaz-K4500= Kavkaz-K4500 L.6.A.4. This experiment was repeated with cultivars Runal, Kavkaz-K4500 L.6.A.4, ST6, TE-9111, Arina, Titlis, M6 and Bulgaria-88 and similar results were obtained.

### The effector cluster resides in a highly dynamic region of the genome

Effector genes are located in plastic, transposable element-rich regions of the genome in many fungal pathogens (Soyer *et al.,* 2014; Dong *et al.,* 2015; Faino *et al.,* 2016). We explored the plasticity of the genomic region harboring the effector gene cluster in order to understand the evolution of *Avr3D1.* With this aim, we performed alignments of the QTL of the 3D7 genome to the genomes of 3D1, the reference strain IPO323 and Swiss strains 1E4 and 1A5. These alignments revealed the absence of *SSP_3* and *SSP_4* in 3D1, IPO323 and 1E4 and the absence of *SSP_3* in 1A5 (Fig. 3a, S9). In order to gain further insight into the plasticity of this effector cluster, we extended our analysis using Illumina genome sequences of 128 *Z. tritici* strains obtained from four different field populations located on three continents (Hartmann & Croll, 2017; Hartmann *et al.,* 2017). *SSP_3* and *SSP_4* were absent in 65% and 42% of the strains, respectively, whereas *Avr3D1* and *QTL7_5* were present in all or 95% of the strains, respectively (Fig. 3b). The presence/absence polymorphisms exhibited by several SSP-encoding genes in this cluster highlight the dynamic nature of the genomic region harboring the virulence QTL.

**Fig. 3.**
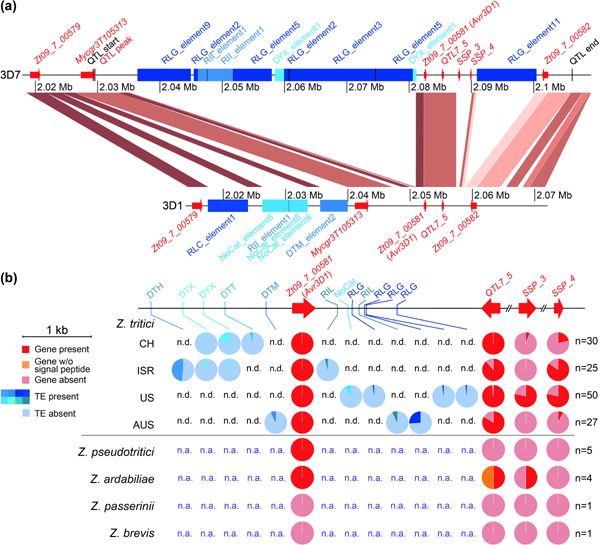
A dynamic and effector-rich region on chromosome 7 is associated with quantitative virulence. (a) Synteny plot comparing the QTL for virulence between strains 3D7 and 3D1. The borders of the 95% confidence interval of the QTL in 3D7 are marked by black vertical lines. Genes are represented by red arrows and transposable elements (TEs) are represented by blue blocks. Collinear sequences between the two strains are shown in different shades of brown indicating sequence identity.(b) Presence/absence polymorphisms of genes predicted to encode small secreted proteins (SSPs) and of TEs in different populations of *Z. tritici* and in four closely related species. The pie charts in red shades show the presence/absence polymorphisms of genes encoding SSPs. The blue pie charts display the presence/absence polymorphisms of TEs in *Z. tritici* strains up-and downstream of the gene *Zt09_7_00581*. The type of TE is indicated, n.d.= not detected, n.a.= not analyzed, n = number of strains.

To investigate whether *Avr3D1*, *SSP3, SSP4* and *QTL7_5* originated after speciation, we analyzed *Z. tritici* sister species to determine if they contained homologs of the genes. A homolog of *Avr3D1* was identified in all examined strains of *Zymoseptoria pseudotritici* and *Zymoseptoria ardabiliae,* but not in *Zymoseptoria brevis* or *Zymoseptoria passerinii,* suggesting that *Avr3D1* originated before *Z. tritici* speciation. Homologs of *QTL7_5* and *SSP_3* were found in only 2 out of 4 strains of *Z. ardabiliae* but not in *Z. pseudotritici* (Fig. 3b). Homologs of *SSP_4* were not identified in the other *Zymoseptoria* species, indicating that this gene may have originated after *Z. tritici* speciation.

We extended our investigation on the genomic plasticity of the effector gene cluster to consider the presence of repetitive elements and TEs. Two insertions of TEs (of 44.5 kb and 9.5 kb, respectively) flanked the four SSP-encoding genes in 3D7, but not in 3D1, where a different TE insertion was present upstream of the QTL (Fig. 2a). The insertion upstream of the SSP genes in 3D7 consisted of an island of 10 different TEs, located 1.3 kb upstream of the start codon of *Avr3D1.* The closest TE to *Avr3D1* is a DNA TE from the Crypton superfamily, which is relatively rare in *Z. tritici.* Upstream of the Crypton element, three different long terminal repeats (LTRs) from the superfamily Gypsy, the most frequent retrotransposons in *Z. tritici* (Grandaubert *et al.,* 2015), were inserted. A Gypsy LTR was also inserted only in 3D7 1 kb downstream of the effector cluster. No TE insertions occurred in the QTL region of the reference strain IPO323 or the Swiss strains 1E4 and 1A5 (Fig. S9). Like in 3D1, TE insertions upstream of the QTL were identified in 1E4 and 1A5 (Fig. S9). Although all the insertions were upstream of the gene *Zt09_7_00580,* they were located at different positions and classified as different superfamilies (Copia in 3D1 and Mutator in 1E4 and 1A5). We extended the analysis of chromosomal rearrangements to the 132 global strains. Remarkably, we observed that 18% of these strains contained at least one TE within 6.5 kb upstream of the cluster. Furthermore, seven different insertions were identified between *Avr3D1* and *QTL7_5.* The inserted TEs belonged to different superfamilies and were located at various positions (Fig. 3b), suggesting that several different insertion events occurred independently. Thus, the effector cluster resides in a highly dynamic region of the genome, in accordance with what has been previously described for other pathogenic fungi in which effectors reside in fast-evolving regions of their two-speed genome (Raffaele & Kamoun, 2012).

### *Avr3D1* is highly polymorphic in four global *Z. tritici* field populations

Escape from recognition is often mediated by modifications in avirulence gene sequences. Therefore, we explored sequence polymorphisms of the avirulence gene *Avr3D1.* In the strain 3D1, the avirulent allele of *Avr3D1 (Avr3D1_3D1_)* encodes a protein of 92 amino acids with a predicted signal peptide of 21 amino acids and a high number of cysteines (8 residues, 11.3%). *Avr3D1* has three exons, of which only exon 1 and exon 2 contain coding DNA. The sequence polymorphism of *Avr3D1* was analyzed in the same four global *Z. tritici* populations used for TE presence/absence analyses. Among these 132 strains, 31 different alleles were identified, encoding 30 different protein variants, all of which were population-specific (Fig. 4a). Strikingly, the 500 bp upstream of the start codon and the 500 bp following the stop codon showed lower diversity (**π**_UP flanking_ = 0.0179; **π**_Down flanking_ = 0.0023) than the coding DNA sequence (CDS; TTCDS = 0.067). In addition, nucleotide diversity was much lower in the first intron (TT¡ntron1 = 0.0003) and the signal peptide sequence (iTsp = 0.0112) compared to the sequence encoding the mature protein (limature protein = 0.068, Fig. 4, S10). This pattern is consistent with accelerated diversification of the CDS, as confirmed by the high ratio between nonsynonymous and synonymous mutations (dN/dS) in the populations (Notes S1). According to the codon-based maximum likelihood approach, 58 out of 96 codon sites were estimated to be under purifying selection, 3 were neutral, and 35 were under diversifying selection (Fig. 4b, Notes S1), suggesting that strong diversifying selection has led to high sequence polymorphism of *Avr3D1.* We hypothesize that numerous adaptive mutations have occurred in this avirulence gene, most probably to counteract recognition by a resistance protein.

**Fig. 4.**
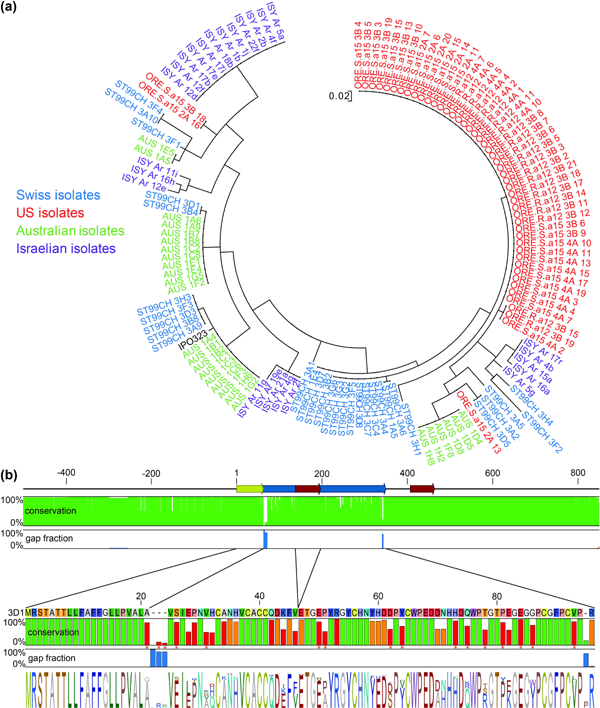
Avr3D1 is highly polymorphic and exhibits the signature of diversifying selection. (a) Phylogenetic tree of the protein sequence of Avr3D1 generated from 132 *Z. tritici* strains from four populations and the reference strain IPO323. (b) Upper panel: Representation of the *Avr3D1* gene including two introns (red arrows), the coding DNA sequence (CDS) of the mature protein (blue arrow) and the signal peptide (yellow arrow). Conservation and gap fractions of each nucleotide in 132 global *Z. tritici* strains is shown for each nucleotide as green and blue vertical bars, respectively. Lower panel: Protein sequence encoded by the avirulent allele *Avr3Dl3Di.* Conservation is shown as vertical bars in green, orange and red, representing purifying, neutral or diversifying selection, respectively, as determined by analysis of dN/dS ratios. Residues under significant (p<0.01) diversifying selection are labelled with red asterisks. Gap fractions are shown as blue vertical bars. The consensus sequence and sequence diversity are depicted as sequence logos.

Despite the high protein diversity, the amino acid substitutions did not affect the signal peptide and did not occur in any of the eight cysteine residues, indicating that the overall backbone structure of Avr3D1 is conserved. Remarkably, in the orthologs in *Z. pseudotritici* and *Z. ardabiliae* (with 60.2% and 53.5% protein identity, respectively) all the cysteine residues were also conserved (Fig. 4, S11, S12). This conservation of the overall protein structure may indicate a general role for Avr3D1 in host colonization that was preserved after speciation.

### Substitutions in *Avr3D1* lead to evasion of recognition

The Avr3D1 variants in the avirulent strain 3D1 and in the virulent strain 3D7 share 86% sequence identity as a consequence of 12 amino acid substitutions and one gap in 3D7 (Fig. 1b). To determine the impact of these differences on recognition, we ectopically expressed the 3D1 *(Avr3D1soi)* and the 3D7 *(Avr3D1_3_D7)* alleles of *Avr3D1* under the control of the promoter from 3D1 in the knockout background and tested the ability to complement the phenotype of 3D1Aavr3D1. *Avr3D1_3D1_* fully complemented the virulence phenotype of 3D1Aavr3D1 with respect to both lesion development and pycnidia production. However, *Avr3D1_3D7_* did not alter the phenotype of 3D1Aavr3D1, indicating that Avr3D1_3D1_ but not Avr3D1_3D7_ triggers an immune response in cv. Runal (Fig. 1a). Moreover, expression of *Avr3D1_3D1_* under the control of the promoter from 3D1 in the 3D7Aavr3D1 background led to a significant reduction in disease (avirulence), whereas expression of *Avr3D1_3D7_* did not alter the phenotype in the same genetic background (Fig. 1a). Therefore, Avr3D13D1, but not Avr3D13D7, is recognized in both genetic backgrounds, demonstrating that substitutions in Avr3D1 led to evasion of recognition in the virulent strain 3D7.

## Discussion

In numerous plant pathosystems, a key determinant of host specificity is the resistance protein-mediated recognition of avirulence factors, which are often SSPs. Though 20 race-specific large-effect resistance genes against *Z. tritici* have been mapped in the wheat genome, their fungal interactors remain unknown with the exception of the resistance gene *Stb6.* Here, we report the discovery of a new *Z. tritici* gene, *Avr3D1,* encoding a cysteine-rich small secreted protein that triggers quantitative resistance in wheat cultivars harbouring the *Stb7* locus.

### Avr3D1 recognition induces quantitative resistance

Avr3D1 is a candidate effector that is expressed during the latent phase but downregulated upon the onset of the necrotrophic phase, suggesting a function in host colonization during the latent phase. The recognition of the avirulent allele leads to a dramatic reduction in the amount of infection and pycnidia formation. This demonstrates that Avr3D1 is an avirulence factor that is likely to be specifically recognized by an *Stb* gene. The fact that only certain wheat cultivars recognize Avr3D1 suggests that recognition follows the gene-for-gene model. In contrast to what has been shown for most other avirulence factors, Avr3D1 recognition does not lead to full resistance, but instead to quantitative resistance in which the pathogen is impaired in its ability to infect, but eventually completes its life cycle. The mechanisms through which *Z. tritici* eventually circumvents the resistance response remain unknown. We hypothesize that the magnitude of the defence response is not strong enough to prevent the progression of the infection and/or that the downregulation of *Avr3D1* during the necrotrophic phase substantially decreases the defence response. However, the strength of the response may still be sufficient to limit propagation of the pathogen under field conditions, in which case the underlying resistance gene could be a valuable source of resistance for breeding programs. Pyramiding of *Stb* resistance genes is an objective in several breeding programs because this approach is thought to be an effective and durable strategy to control septoria tritici blotch in the field (Chartrain *et al.,* 2004; Kettles & Kanyuka, 2016). In fact, TE-9111 and Kavkaz-K4500 L6.A.4, two of the cultivars that specifically recognized Avr3D1, contain at least three *Stb* genes and are major sources of resistance to *Z. tritici* (Chartrain *et al.,* 2004). Our work might contribute to the identification of the corresponding *Stb* gene in the future.

In this work, we show that asexual reproduction can occur even upon induction of effector-triggered defence. In the case of AvrStb6 recognition, Stb6 strongly hinders the progression of infection, abolishing the induction of necrosis (Kema, GHJ *et al.,* 2000; Ware, 2006; Zhong *et al.,* 2017). In contrast, recognition of Avr3D1 triggers a weaker form of resistance that prolongs the asymptomatic phase, while allowing necrotic lesions to develop and pycnidia to form. These findings highlight the continuum between qualitative and quantitative resistance in gene-for-gene interactions. Although we identified an avirulence gene that has a large effect on some wheat cultivars, additional factors must contribute to the differences in virulence between the two strains, because the density of pycnidia formed by the Avr3D1 knockout in the avirulent strain was still lower than the pycnidia density produced by the virulent strain. The provided data highlight the quantitative nature of wheat-Z. *tritici* interactions.

### Chromosome rearrangements contribute to diversification of the effector gene cluster

*Avr3D1* and the three other genes in the effector gene cluster are located on the right arm of chromosome 7, which is distinctive because of its low overall expression levels (Rudd *et al.,* 2015) and its enrichment in heterochromatin (Schotanus *et al.,* 2015). In fact, it was postulated that this region originated from a fusion between an accessory chromosome and a core chromosome (Schotanus *et al.,* 2015). Numerous independent insertions of transposable elements surrounding *Avr3D1* were identified in 132 global strains of *Z. tritici.* Transposable elements are frequently described as an evolutionary force shaping adjacent regions by contributing to diversification through non-homologous recombination or RIP (Faino *et al.,* 2016; Wicker *et al.,* 2016). Given that four putative effector genes are clustered in this region, transposable elements could play a similar role in facilitating rapid evolution of these effectors, but may also enable concerted expression of effector genes during infection by chromatin remodelling (Soyer *et al.,* 2014; Schotanus *et al.,* 2015). In the case of this effector gene cluster, transposable elements might have contributed to the high diversity of the avirulence gene *Avr3D1* and the presence/absence polymorphisms shown for the other effector genes. Sequence diversification is particularly relevant for pathogen effectors, as they are key players in the coevolution with their hosts. Indeed, sequence modifications of Avr3D1 in the virulent strain allowed an escape from recognition by the corresponding resistance protein.

### Avr3D1 sequence variation to evade recognition

A common evolutionary strategy for evading recognition is the loss of an entire avirulence gene (Schürch *et al.,* 2004; Mackey & McFall, 2006; de Jonge *et al.,* 2012; Hartmann *et al.,* 2017). However, loss of *Avr3D1* was not observed in any of the 132 global strains, despite its location in a highly plastic genomic region, as shown by presence/absence polymorphisms for neighbouring genes and transposable elements. Other deleterious mutations such as frameshifts, premature stop codons and non-functional splice sites were not found. Despite the high overall diversity, all the cysteine residues and the signal peptide, two core features of effector proteins, were completely conserved. The absence of any high-impact mutations suggests that loss of *Avr3D1* may impose a significant fitness cost. We therefore hypothesize that Avr3D1 plays a crucial role in the life history of *Z. tritici*, though we could not demonstrate a contribution of Avr3D1 to lesion or pycnidia formation in susceptible varieties during the seedling stage under greenhouse conditions. It is possible that the role of Avr3D1 is more pronounced under field conditions or at different developmental stages, e.g. in adult plants. An additional hypothesis to explain the apparent dispensability of Avr3D1 is that functional redundancy masks phenotypic effects in the knockout mutants (Marshall *et al.,* 2011; Win *et al.,* 2012; Mirzadi Gohari *et al.,* 2015; Rudd *et al.,* 2015).

## Conclusion

We identified a new major avirulence factor of *Z. tritici* (Avr3D1) that we hypothesize is recognized by Stb7 or Stb12. Unlike what has been described for most described avirulence factors, recognition of Avr3D1 does not prevent lesion formation or pathogen reproduction, demonstrating that race-specific resistance is not always qualitative. Finally, our comprehensive comparative genomic analysis suggests that effectors in *Z. tritici* are located in dynamic genomic compartments favouring rapid evolution, which may facilitate adaptation to the evolving wheat host.

## Acknowledgments

We thank Zacharie Ashley Ngamenie, Susanne Meile, Anna Spescha and Barryette Oberholzer for their help in various experiments. Eva Stukenbrock and Jason Rudd provided the vectors pES1, pES6 and pCGEN. We thank Marc-Henri Lebrun and Thierry Marcel who provided us with wheat seeds. qPCR was performed in collaboration with the Genetic Diversity Centre (GDC), ETH Zurich. The research was supported by the Swiss National Science Foundation (Grants 31003A_155955, 31003A_173265), ETH Zurich Research Commission Grant 12-03 and INRA Young Scientist grant.

## Author contribution

L.M. and A.S.-V. conceived and designed experiments. L.M., F.H., C.P. and P.B. performed the experiments. L.M., D.C. and A.S.-V. analyzed the data. L.M. and A.S.-V. wrote the manuscript. P.B., D.C., F.H. and B.M. review and edited the manuscript.

## Supporting information

**Fig. S1.** The *Zymoseptoria tritici* strain 3D1 shows lower virulence and a delayed onset of symptoms on cv. Runal.

**Fig. S2.** Manual annotation of putative effector genes in the QTL for virulence.

**Fig. S3.** The gene *QTL7_5* does not contribute to virulence.

**Fig. S4.** *Z09_7_00581* encodes the avirulence factor Avr3D1 and sequence modifications lead to evasion of recognition.

**Fig. S5.** *Avr3D1* does not explain all the differences in virulence between 3D1 and 3D7.

**Fig. S6.** *In vitro* growth of the mutant lines was unaltered under several stress conditions.

**Fig. S7.** *Avr3D1* expression peaks at the end of the latent phase.

**Fig. S8.** Specific recognition of Avr3D1 by certain wheat varieties leads to a reduction in pycnidia formation.

**Fig. S9.** Synteny plot of the QTL between five *Zymoseptoria tritici* strains.

**Fig. S10.** Distribution of nucleotide diversity of *Avr3D1* among *Z. tritici* strains.

**Fig. S11.** Orthologous sequences of Avr3D1 identified in five strains of *Zymoseptoria pseudotritici.*

**Fig. S12.** Orthologous sequences of Avr3D1 identified in four strains of *Zymoseptoria ardabiliae*.

**Fig. S13.** Frequency distribution of dN/dS ratios for all pairwise *Avr3D1* haplotype comparisons.

**Table S1.** Primers used in this study.

**Table S2.** Effector gene cluster annotation. gff file of the manually reannotated effector genes identified in the QTL.

**Table S3.** Model test and parameter estimates of diversifying selection with PAML based on the total *Avr3D1* data set.

**Notes S1.** Population genetic analysis.

